# Addressing the dNTP bottleneck restricting prime editing activity

**DOI:** 10.1101/2023.10.21.563443

**Authors:** Karthikeyan Ponnienselvan, Pengpeng Liu, Thomas Nyalile, Sarah Oikemus, Anya T. Joynt, Karen Kelly, Dongsheng Guo, Zexiang Chen, Jeong Min Lee, Celia A. Schiffer, Charles P. Emerson, Nathan D. Lawson, Jonathan K. Watts, Erik J. Sontheimer, Jeremy Luban, Scot A. Wolfe

**Author notes:** These authors contributed equally: Karthikeyan Ponnienselvan and Pengpeng Liu.

## Abstract

Prime editing efficiency is modest in cells that are quiescent or slowly proliferating where intracellular dNTP levels are tightly regulated. MMLV-reverse transcriptase - the prime editor polymerase subunit - requires high intracellular dNTPs levels for efficient polymerization. We report that prime editing efficiency in primary cells and *in vivo* is increased by mutations that enhance the enzymatic properties of MMLV-reverse transcriptase and can be further complemented by targeting SAMHD1 for degradation.

---

Prime editing, due to its versatility in performing targeted base conversions, deletions and insertions, has the potential for broad therapeutic application in the treatment of a variety of human diseases. A prime editor (PE) has 3 core components: {1} Cas9 nickase, {2} M-MLV reverse transcriptase (MMLV-RT), and {3} prime editing guide RNA (pegRNA), where the pegRNA provides the template for MMLV-RT to introduce sequence changes within the genome through extension of the nicked DNA strand^1,2^. Since the initial description of prime editing^1^, many of the components within the system have been further optimized: including the Cas9 nickase^2^, the nuclear localization sequences^2,3^, pegRNA designs^4–6^ and the DNA repair pathways that are harnessed^2^.

The original PE2 system^1^ utilizes a pentamutant MMLV-RT that increases thermostability, affinity for the RNA-DNA template and processivity. The improved PEmax system also utilizes the pentamutant MMLV-RT^2^. More recently, L139P, a mutation that improves thermostability, affinity for the template, and processivity^7^ was introduced in the pentamutant MMLV to achieve a modest increase in prime editing rates^8^. Other studies have substituted alternate RTs^9–11^, split prime editor systems^9–11^, or utilized molecular evolution to enhance the properties of multiple RTs for prime editing^12^ with notable improvements in precise editing outcomes for some templates. Despite these advances to the prime editing system, precise editing rates remain modest in primary cells or *in vivo* in many tissue types^6,13,14^.

We believe that prime editing efficiency in primary cells is hampered in part by inherent properties of the MMLV-RT, which can potentially be addressed by harnessing the wealth of optimization performed on this enzyme^15^. We observed protein aggregation during purification of the PEmax prime editor protein that is likely a property of the MMLV-RT domain^6^, which suggests that its solubility can be improved. Additionally, most characterization and optimization of prime editing systems has been carried out in rapidly dividing mammalian cells^1,8,10,12^, where high intracellular dNTP levels should facilitate prime editing efficiency^16^. Cellular dNTP levels are 6- to more than 100-fold lower in post-mitotic or quiescent cells than cycling cells^16–19^, which would restrict MMLV-RT activity due to its low affinity for dNTPs^20^. Altering the conserved Y**<u>V</u>**DD motif in MMLV-RT to Y**<u>M</u>**DD (V223M) changes its active site from an onco-retroviral RT to a more efficient lentiviral-like RT, reducing its *K_m_* for dNTPs by 2 to 4-fold^21–23^. Consequently, we assessed the impact of the V223M mutation and the L435K mutation, which increases MMLV-RT solubility^24^, on prime editing activity.

We introduced the L435K and V223M mutations in our PEmax protein expression construct, which contains four nuclear localization signal sequences^6^, to evaluate their impact on protein solubility and prime editing activity (designated PEmax**, Extended Data Fig. 1a). Consistent with our prior experience, the purified PEmax protein could not be concentrated above 30 μM without aggregate formation. In contrast, the purified PEmax** protein was concentrated to 140 μM without visible aggregates. The improved solubility of the PEmax** prime editor protein resulted in a 7-fold increase in protein yield (Extended Data Fig. 1b,c). We evaluated the prime editing efficiency of PEmax and PEmax** proteins under limiting dNTP concentrations. The incorporation rate of a 20-nucleotide sequence tag^25^ rose with increasing dNTP concentration with a plateau at ∼0.2 μM dNTPs for both proteins (Figure 1a). PEmax** displayed significantly higher enzymatic activity than PEmax when dNTP levels were subsaturating (0.05 to 0.15 μM dNTPs), consistent with the improved dNTP affinity of the V223M mutation^22^.

**Figure 1:**
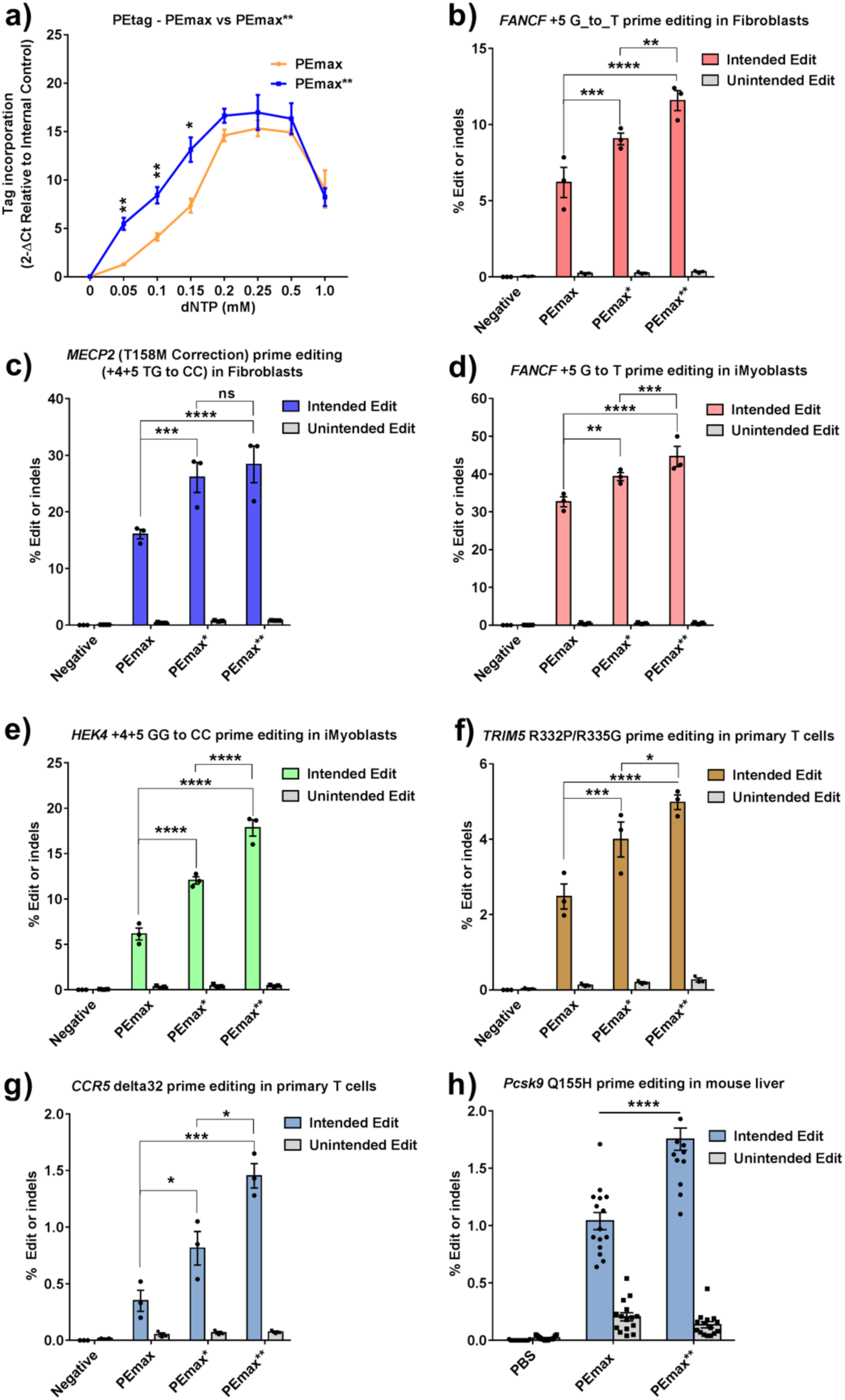
PEmax** increases prime editing rates across different cell types and in vivo. (**a**) *In vitro* biochemical analysis of PEmax and PEmax** RNP activity under different dNTP concentrations assessing 20 nt tag incorporation at the HEK4 target site in purified gDNA (qPCR, n=3 independent replicates). Two-way ANOVA was used to compare PEmax and PEmax** for each condition; * indicates P ≤0.05, ** indicates P ≤0.01 (also see Supplementary table). (**b-g**) PEmax, PEmax* (V223M) and PEmax** (V223M & L435K) mRNA mediated prime editing efficiency with synthetic pegRNA/sgRNA delivered by electroporation in a variety of cell types: (**b**) PE3 editing at *FANCF* (+5 G to T) and (**c**) PE2 editing at *MECP2* (+4+5 TG to CC) to revert the T158M mutation in patient fibroblasts (n=3 independent replicates), (**d**) PE3 editing at *FANCF* (+5 G to T) and (**e**) HEK4 (+4+5 GG to CC) in induced myoblasts (n=3 independent replicates), and PE3 editing to install HIV-1 resistance mutations (**f**) *CCR5^delta^*^32^ and (**g**) *TRIM5ɑ*^R332P,R335G^ in activated primary human T cells (n=2 different donors). (**h**) Comparison of PE3 editing rates *in vivo* at *Pcsk9* (Q155H) mediated by PEmax and PEmax** mRNA LNP formulations delivered to mice through retro-orbital injection (rates determined from 3 liver punches from each of 5 mice in each cohort, n = 15). Two-way ANOVA was used to compare the intended edit and unintended edit across all the groups for each graph, PEmax was used as a control column for multiple comparisons; ns indicates P > 0.05, * indicates P ≤0.05, ** indicates P ≤0.01, *** indicates P ≤0.001 and **** indicates P ≤0.0001 (also see Supplementary table).

Next, we screened the impact of PEmax^V223M^ (PEmax*), PEmax^L435K^ and the double mutant PEmax^V223M,L435K^ (PEmax**) on prime editing activity in HEK293T cells and U2OS cells. We performed prime editing under subsaturating conditions for plasmid delivery by transient transfection to discern differences in activity. We utilized pegRNAs designed to insert a 20 nucleotide sequence to assess RT processivity at two well-characterized target sites: HEK4 and *VEGFA* (Extended Data Fig. 2)^25^. In both HEK293T and U2OS cells across both target sites, we observed a ∼2-fold increase in precise editing rates for PEmax^L435K^ compared to PEmax. In contrast, we observed no significant change in precise editing rates for PEmax*. Consistent with the single-mutant analysis, we observed a ∼2-fold increase in precise editing rates for PEmax** suggesting that the L435K is the primary driver of improved prime editing activity in transformed cell lines that are rapidly dividing.

We compared the *in vivo* prime editing activity of PEmax and PEmax** ribonucleoprotein (RNP) complexes in zebrafish embryos. We designed PE RNPs programmed with a synthetic pegRNA (PE2 format) to generate two different non-synonymous mutations - R841W or L907F - within the *tek* locus^6^. Encouragingly, we observed a ∼2-fold increase in precise editing rates for PEmax** RNPs for introduction of the R841W mutation. Dose-dependent editing was observed, where precise editing rates reached ∼30% (Extended Data Fig. 3a). A similar result was observed for introduction of the L907F mutation. PEmax** RNPs produced ∼2-fold improvement in genome editing, where precise editing rates reached ∼30% (Extended Data Fig. 3b).

We also evaluated prime editing activity in induced myoblasts (iMyoblasts), which are differentiated from iPSCs using a transgene-free myogenic protocol^26^. We compared PEmax or PEmax** RNPs programmed with a synthetic pegRNA and a nicking sgRNA targeting the *FANCF* or HEK4 target sites (PE3 format), two benchmark sites that are commonly employed for assessing prime editing efficiency^1^. Similar to the observations in zebrafish embryos, we observed a 1.3- to 1.7-fold increase in precise editing outcomes when employing PEmax** RNPs with maximal editing efficiencies ranging from 15 to 40% (Extended Data Fig. 3c,d).

After demonstrating the increased precise editing rates for PEmax** RNPs in mammalian stem cells and *in vivo*, we compared the activity of PEmax, PEmax^V223M^ (PEmax*) and PEmax^V223M,L435K^ (PEmax**) in multiple different cell types to dissect the contribution of each mutation to prime editing. In these experiments we co-delivered *in vitro* transcribed mRNA encoding each prime editor with synthetic pegRNA and a nicking sgRNA when employing PE3 editing. We evaluated prime editing in a patient-derived fibroblast line carrying a common mutation associated with Rett syndrome (T158M in *MECP2*)^27^. We observed a 1.5- and 1.9-fold increase in prime editing at *FANCF* with PEmax* and PEmax** (PE3), respectively, compared to PEmax (Figure 1b). Likewise, PEmax* and PEmax** (PE2) increased the correction rate of the *MECP2* T158M mutation by approximately 1.6- and 1.8-fold, respectively (Figure 1c). We also evaluated prime editing efficiency in iMyoblasts. We observed a 1.2-fold and 1.4 fold increase in prime editing at *FANCF* with PEmax* and PEmax**, respectively (Figure 1d). We observed a 2.0-fold and 2.9-fold increase in prime editing at HEK4 with PEmax* and PEmax**, respectively (Figure 1e). These data indicate that introduction of the V223M mutation (PEmax*) is beneficial for increasing precise editing outcomes in primary cell types and the inclusion of the L435K mutation (PEmax**) further improves prime editing rates.

To explore the utility of PEmax variants for prime editing in other therapeutically relevant cell types, we tested their ability to introduce HIV-1 resistance mutations in activated human primary T cells. We used prime editing to introduce R332P and R335G mutations in *TRIM5α* that restrict HIV-1 infection^28^. We observed a 1.6-fold increase in precise editing rates with PEmax* and 1.9-fold increase with PEmax** when compared with PEmax (Figure 1f). Next, we introduced the delta32 mutation within *CCR5* that is associated with HIV-1 resistance^29^. We observed a 2.2-fold increase with PEmax* and 3.8-fold increase with PEmax** in the precise 32 bp deletion when compared with PEmax (Figure 1g). A similar improvement in the precise introduction of *CCR5^delta^*^32^ (5.3 fold) was observed for PEmax** in resting T cells, although the overall editing rate was modest (∼0.3%, Extended Data Fig. 4).

Motivated by the observed improvements in prime editing with PEmax** in multiple cell types, we tested its activity in mice. We introduced the disabling Q155H mutation in *Pcsk9* recently installed by the Liu lab using a split-intein prime editing system delivered by dual adeno-associated virus (AAV)^14^. We delivered LNPs by retro-orbital injection in young adult mice containing PEmax or PEmax** mRNA along with the optimized synthetic epegRNA and nicking guide RNA described in their study^14^. We observed 1.8% precise editing at *Pcsk9* with PEmax**, a 1.7-fold improvement over PEmax editing in the mouse liver (Figure 1h). Thus, PEmax** increases precise editing rates *in vivo* in adult animals.

Our biochemical and cellular observations are consistent with intracellular dNTP levels limiting prime editing efficiency in slowly proliferating or quiescent cells. Cells employ multiple pathways to regulate dNTP levels to control unplanned DNA synthesis^30^. SAMHD1 is a deoxynucleotide triphosphohydrolase that maintains low dNTP levels in cells that are not undergoing DNA replication, which restricts infection by retroviruses^31–33^. Lentiviruses like HIV-2 package an accessory protein - VPX - that targets SAMHD1 for degradation via the cullin 4-based (CRL4) E3 ubiquitin ligase pathway^34,35^. SAMHD1 proteolysis increases cellular dNTP concentration, which promotes reverse transcription and successful viral infection in quiescent cells^32,33^. Therefore, we investigated if co-delivering VPX would increase prime editing rates in resting and activated T cells. Co-delivery of VPX protein increased precise editing rates for PEmax, PEmax* or PEmax** at the *FANCF* locus (Extended Data Fig. 5a,b). Co-delivery of increasing amounts of VPX protein progressively increased precise editing rates in resting T cells without reaching saturation within our titration range (Extended Data Fig. 5c). Co-delivery of VPX protein with PEmax** increased the introduction of the *TRIM5ɑ* double mutation in activated T cells (Extended data 5d). Next, we compared the efficiency of PE3 editing with PEmax to the TWIN-PE approach^36^ with Cas9-NG^37^ PEmax for introduction of *CCR5^delta^*^32^ in resting T cells^37^. We observed no substantial difference between a pegRNA and epegRNA for PE3 editing, but TWIN-PE introduced the *CCR5^delta^*^32^ mutation at significantly higher rates (Extended Data Fig. 5e). The TWIN-PE approach for introduction of *CCR5^delta^*^32^ was significantly more efficient with co-delivered VPX protein in activated T cells (Extended Data Fig. 5f).

We also produced VPX and VPX^Q76A^ mRNA for co-delivery with the PEmax variants. VPX^Q76A^ cannot interact with DCAF1 (the E3 ubiquitin ligase substrate adaptor^34^), and consequently cannot target SAMHD1 for proteasomal degradation. We observed a stimulatory effect on prime editing rates when the VPX mRNA was co-delivered in patient derived fibroblasts and resting T cells, whereas no improvement was observed when VPX^Q76A^ mRNA was co-delivered (Figure 2a,b). VPX mRNA also stimulated prime editing in activated T cells at therapeutically relevant targets. We co-delivered PEmax** mRNA and an epegRNA for introducing the *TRIM5ɑ* double mutation with VPX protein and observed a 1.5-fold increase in editing (Figure 2c). Likewise, co-delivery of VPX mRNA increased the efficiency for the *CCR5^delta^*^32^ TWIN-PE strategy by 1.5-fold with application in one donor achieving >40% precise edits (Figure 2d). Thus, VPX delivered as protein or mRNA can provide a potent boost to prime editing rates in a variety of primary cell types.

**Figure 2:**
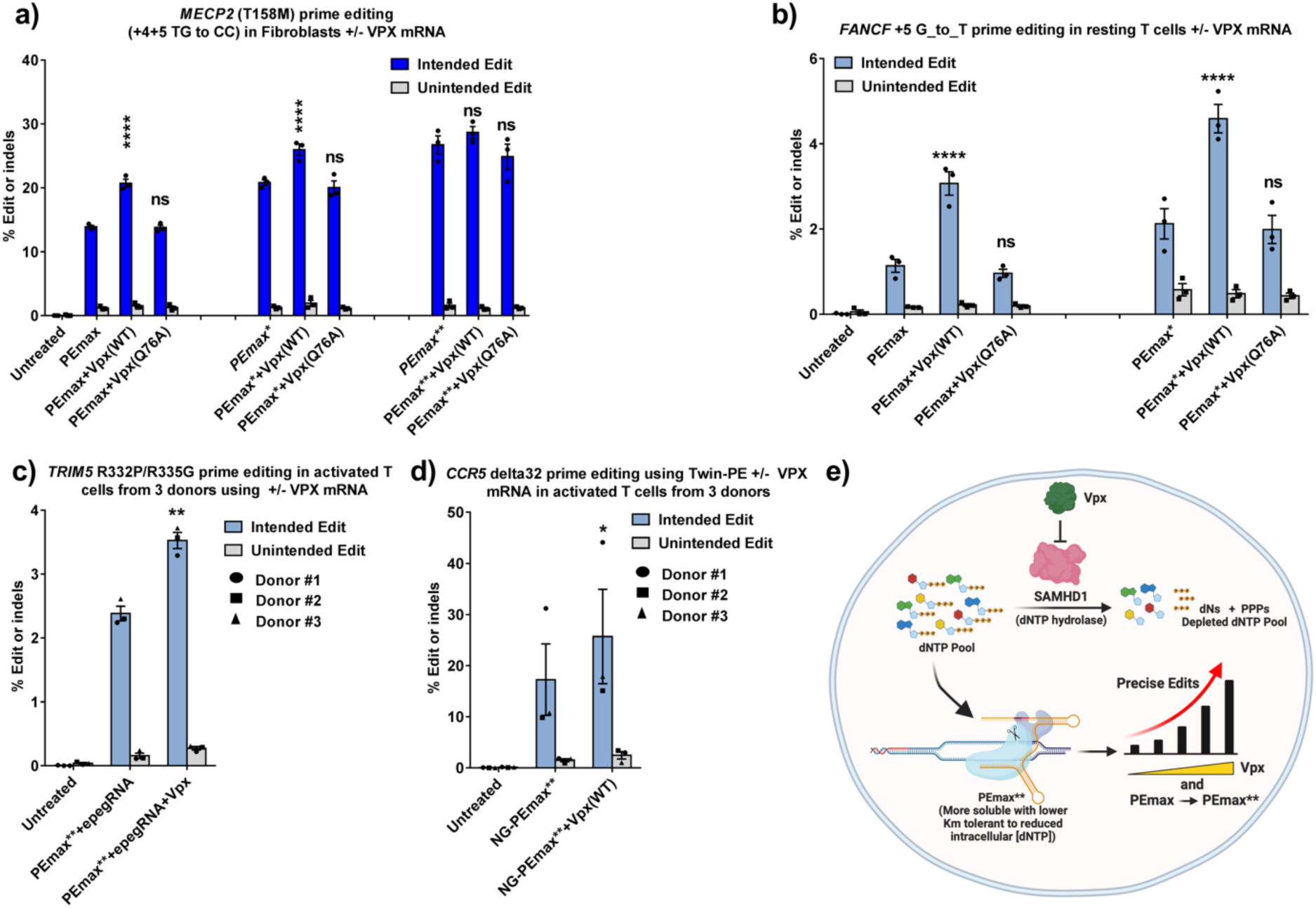
Co-delivery of VPX mRNA increases prime editing rates in patient-derived fibroblasts and human primary T cells. **(a)** PEmax, PEmax* (V223M) and PEmax** (V223M & L435K) mRNA-mediated PE2 editing at *MECP2* (+4+5 TG to CC) to revert the T158M mutation in patient derived fibroblasts with co-delivery of synthetic pegRNA and VPX or VPX^Q76A^ mRNA (n=3 independent replicates). PEmax/PEmax*/PEmax** were individually used as a control column for statistical comparisons within three groups. **(b)** PEmax and PEmax* mRNA-mediated PE3 editing at *FANCF* (+5 G to T) in resting human primary T cells with co-delivery of synthetic pegRNA/sgRNA and VPX or VPX^Q76A^ mRNA (n=3 different donors). PEmax and PEmax* were used as a control column for statistical comparisons within these two groups. (**c**) PE3 editing to install *TRIM5ɑ*^R332P,R335G^ HIV-1 resistance mutations in activated human primary T cells using PEmax** and a synthetic epegRNA with or without co-delivery of VPX mRNA (n=3 different donors). (**d**) Twin-PE editing to install *CCR5^delta^*^32^ HIV-1 resistance mutation in activated primary human T cells using PEmax** with Cas9-NG PAM preference with or without co-delivery of VPX mRNA (n=3 different donors). Two-way ANOVA was used to compare the intended edit and unintended edit across all the groups for each graph, PEmax** or NG-PEmax** was used as a control column for multiple comparisons in (c) and (d), respectively; ns indicates P > 0.05, * indicates P ≤0.05, ** indicates P ≤0.01, *** indicates P ≤0.001 and **** indicates P ≤0.0001 (also see Supplementary table). (**e**) Schematic overview of methods (PEmax** and VPX) used to increase editing rates in cell populations with restricted dNTP levels.

In this study, we have identified two bottlenecks (solubility and dNTP affinity) that restrict the activity of MMLV-RT-based prime editing systems in primary cells and *in vivo*. We introduced L435K and V223M point mutations into the MMLV-RT to improve solubility and reverse transcription activity at low dNTP concentrations, respectively^22,24^ (Supplementary Note). Improved solubility translated to improved prime editing for every tested delivery modality (protein, mRNA and plasmid). The apparent dNTP restriction to prime editing observed in primary cells and *in vivo* is consistent with previous observations of retroviral infection of resting cells where increased dNTP levels (achieved through either extracellular deoxynucleoside supplementation or SAMHD1 knockdown) improve infection rates^32,33,38,39^. Interestingly, no benefit was observed in rapidly dividing transformed cell lines for the V223M mutation. The effect of the dNTP mutant was more prominent in resting T cells, which have lower levels of dNTPs^17^. Given the range of applications where PEmax** improves editing rates, we believe that this variant will have broad utility in editing quiescent and post-mitotic cells. In addition, PEmax** protein provides high rates of precise genome editing in zebrafish embryos, which may facilitate the analysis of mutant phenotypes in the F0 generation.

While this study was being submitted, the Liu laboratory described the development of a suite of compact prime editor variants evolved from their PANCE and PACE selection systems^12^. These RT variants were selected for improved activity and processivity in bacteria, which translated to improved prime editing with longer, structured pegRNA templates in mammalian cells. Their PE6d variant, which employs a truncated MMLV-RT lacking the RNaseH domain, includes a V223Y mutation, however it is unclear if that mutation improves dNTP affinity like V223M. The PE6d variant displayed improved processivity, which is evident by the increased proportion of pegRNA scaffold sequence incorporation within the unwanted editing products. We observe a similar increase in the proportion of pegRNA scaffold sequence incorporation within the unwanted editing products for PEmax* and PEmax** (Extended Data Figure 6a-d). Similar to V223M, the V223A mutation in the MMLV-RT improves its dNTP affinity^22^ and inclusion of this mutation in a recently engineered plant prime editor increased prime editing rates^40^. DNA polymerase-based prime editing systems may also provide avenues to overcome low dNTP levels that restrict prime editing in quiescent and post-mitotic cells due to their superior processivity and dNTP affinity relative to RTs^41,42^. However, innate immune sensing of their DNA templates within the cytoplasm may present an obstacle to editing in some systems^43^.

To further facilitate MMLV-RT processivity, we knocked down SAMHD1, which is involved in maintaining low intracellular levels of dNTPs^31^. For that purpose, we utilized VPX, a viral accessory protein that targets SAMHD1 for proteasome-mediated degradation^32^. We show that VPX protein or mRNA can be co-delivered along with the prime editing reagents to increase editing rates in patient-derived fibroblasts and primary human T cells. Using PEmax** with co-delivered VPX increased prime editing rates up to 4-fold over PEmax alone. SAMHD1 also plays a role in DNA resection and repair at collapsed replication forks^44,45^. However, we did not observe an obvious impact of VPX co-delivery on the distribution of indels that are produced by prime editing (Extended Data Fig. 7a-d). We envision that VPX co-delivery with our improved prime editing reagents may be used with LNPs or eVLPs^46^ to improve editing outcomes in various organ systems *in vivo* (Fig. 2e).

## Acknowledgements

We thank members of the Emerson, Lawson, Luban, Schiffer, Sontheimer, Watts and Wolfe labs for helpful discussions. Rett Syndrome Research Trust for providing Rett fibroblasts. J.M.L. and C.A.S. were supported by a grant from the National Institutes of Health (R01AI150478). D.G. and C.P.E.Jr were supported by a grant from the National Institutes of Health (U54HD0060848). S.O. and N.D.L. were supported by grants from the National Institutes of Health (R35HL140017 and R21OD030004). Z.C. and E.J.S. were supported by a grant from the National Institutes of Health (R01GM150273) and the Leducq Foundation. K.K., and J.W. were supported in part by the National Institutes of Health (UG3TR002668). T.N. and J.L. were supported by a grant from the National Institutes of Health (R37AI147868 & U54AI170856). K.P., P.L., A.T.J. and S.A.W. were supported in part by the National Institutes of Health (R01HL120669, R37AI147868 and UG3TR002668), the Doris Duke Charitable Foundation, and the Rett Syndrome Research Trust.

## Author contributions

K.P., P.L., J.L. and S.A.W. conceptualized the project, K.P., P.L., Z.C., C.P.E.Jr, N.D.L., E.J.S., J.W., J.L. and S.A.W. designed experiments. K.P., P.L., T.N., A.T.J., D.G. and Z.C. conducted molecular biological and cellular experiments. K.P., P.L. and J.M.L. purified prime editor proteins. K.P., P.L., and S.O. conducted zebrafish editing experiments. K.P., P.L., A.T.J., Z.C. and K.K. conducted mouse editing experiments. K.P. and P.L. conducted high-throughput sequencing and bioinformatic analyses. K.P., P.L., T.N., J.L. and S.A.W. interpreted the data and wrote the paper, and all authors edited the paper.

## Materials and methods

### General methods and molecular cloning

Expression plasmids used for pegRNAs have been previously described^47^. Mammalian expression plasmids for different PEmax variants were generated by site-directed mutagenesis on pCMV-PEmax (a gift from David Liu, Addgene plasmid #174820). All plasmids used for transient transfection experiments were purified with an endotoxin removal step (ZymoPURE Plasmid Miniprep Kit from Zymo Research). The PEmax** protein expression vector (pET-21a-PEmax**-6His) was generated by amplifying the fragment of MMLV-RT containing the V223M and L435K mutations from pCMV-PEmax**. This PCR product was digested with BamHI and BsrGI and cloned into pET-21a-PEmax-6His (Addgene plasmid #204471) digested with the same enzymes. The PEmax* and PEmax** mRNA *in vitro* transcription vector was generated by amplifying the fragment of MMLV-RT mutants from pCMV-PEmax* and pCMV-PEmax** separately (Supplementary Figure 1). Then PCR products were Gibson-cloned into the PEmax mRNA plasmid (Addgene plasmid #204472) digested with RsrII and XhoI enzymes. VPX coding sequence was amplified from pscALPS gag-gfp/deltavpx vector (Addgene #115807, a gift from Jeremy Luban) and inserted into PEmax mRNA plasmid (Addgene plasmid #204472) digested by SalI and EcoRI. The VPX^Q76A^ vector was generated from the VPX mRNA vector by Gibson cloning.

### pegRNA designs

We designed the pegRNAs with a primer binding site (PBS) length and composition that reduces the auto-inhibitory interaction between the PBS and spacer sequence within the pegRNA, as described in our previous study^6^, we utilized the MELTING 5 program^48^ to identify a PBS length nearest to the ideal T_m_ value identified (37°C). The RNA sequence of the PBS was entered in the 5’ to 3’ direction along with the nucleic acid concentration, hybridization type and sodium concentration based on the 10 µL RNP complex reaction conditions (nucleic.acid.concentration = 2e-05, hybridization.type = “rnadna”, Na.conc = 1 and method.nn = “sug95”). Based on the output of predicted T_m_ values, a PBS sequence length was chosen that has a T_m_ close to 37°C. The PBS length for all pegRNAs used in this study except the *PCSK9* pegRNA were designed using the MELTING 5 program. For epegRNAs design, we used pegLIT (<u>peglit.liugroup.us</u>) to identify a suitable linker sequence by inputting the spacer, PBS, and reverse transcriptase template (RTT) for each specific target site^4^.

### In vitro transcription of mRNA for PEmax, PEmax variants and VPX

PEmax, PEmax*, PEmax**, VPX and VPX^Q76A^mRNA plasmid were linearized by the PmeI enzyme that cleaves after the polyA tail. mRNA was transcribed from 500 ng purified linearized template using the HiScribe T7 High-Yield RNA Synthesis Kit (New England BioLabs) with co-transcriptional capping by CleanCap AG (TriLink Biotechnologies) and full replacement of UTP with N1-Methylpseudouridine-5’-triphosphate (TriLink Biotechnologies). After 1 hour of *in vitro* transcription, the DNA template was digested by 1 μL DNaseI (Thermo Fisher Scientific) for 15 min. Transcribed mRNAs were purified by RNA Clean & Concentrator-25 kit from Zymo Research, then purified mRNA was dissolved in nuclease-free water. The resulting PEmax mRNA was quantified with a NanoDrop One UV-Vis spectrophotometer (Thermo Fisher Scientific) and stored at −80°C.

### PEmax and PEmax** protein purification

PEmax protein purification followed our previously described protocol^6^. Briefly, PEmax and PEmax** protein expression constructs were introduced into *E. coli* Rosetta2(DE3)pLysS cells (EMD Millipore). Bacteria were grown at 37°C in baffle flasks to an OD600 of ∼0.6, then pre-chilled in an ice bath for 10 minutes and shifted to growth at 18°C. At an OD600 of ∼0.8 the cells were induced for 16 hours with IPTG (0.7 mM final concentration). Following induction, cells were pelleted by centrifugation (3500 g, 20 min) and then resuspended with Nickel-NTA buffer (20 mM TRIS + 1 M NaCl + 20 mM imidazole + 1 mM TCEP, pH 7.5) supplemented with HALT Protease Inhibitor Cocktail, EDTA-Free (100X) [ThermoFisher] and lysed with LM-20 Microfluidizer (Microfluidics) following the manufacturer’s instructions. The lysate was transferred to a centrifuge tube and spun at 20,000 g for 20 minutes. The clarified lysate was then purified with Ni-NTA resin (Qiagen) in batch mode, washed with wash buffer (20 mM TRIS + 1 M NaCl + 20 mM imidazole + 1 mM TCEP, pH 7.5) and eluted with an elution buffer (20 mM TRIS, 500 mM NaCl, 250 mM Imidazole, 10% w/v glycerol, pH 7.5). The eluted proteins were dialyzed overnight at 4°C in 20 mM HEPES, 500 mM NaCl, 1 mM EDTA, 10% w/v (8% v/v) glycerol, pH 7.5. Subsequently, the proteins were step-dialyzed from 500 mM NaCl to 250 mM NaCl to 200 mM NaCl (not exceeding one hour incubation per step; final dialysis buffer: 20 mM HEPES, 200 mM NaCl, 1 mM EDTA, 10% w/v glycerol, pH 7.5). Next, the proteins were purified by anion and cation exchange chromatography [the protein was loaded on a stacked column of 5ml HiTrap-Q HP and 5ml HiTrap-S (Cytiva), Buffer A = 20 mM HEPES pH 7.5 + 1 mM TCEP, Buffer B = 20 mM HEPES pH 7.5 + 1 M NaCl + 1 mM TCEP, Flow rate = 5 ml/min, CV = column volume = 5 ml]. The anion exchange column was removed prior to the elution of the prime editor protein from the cation exchange column. The primary prime editor protein peak fractions were dialyzed into 20 mM HEPES pH 7.5, 300 mM NaCl and then concentrated to ∼30μM for PEmax and 140μM for PEmax** using a 100 kDa amicon ultra centrifugal filter (UFC910008, Millipore).

### Culture conditions for immortalized cell lines and patient-derived fibroblasts

HEK293T cells and U2OS cells were purchased from ATCC. Patient-derived fibroblasts containing the T158M mutation were a gift from the Rett Syndrome Research Trust. All cells were maintained in Dulbecco’s Modified Eagle’s Medium supplemented with 10% FBS at 37°C and 5% CO_2_.

### Transfection of HEK293T and U2OS cells

HEK293T and U2OS cells were plated 40,000 cells per well in a 48-well plate. 24 hours later, the cells were co-transfected with 200 ng of prime editor plasmid and 100 ng of pegRNA plasmid using Lipofectamine 3000 (Invitrogen) according to the manufacturer’s instructions. To determine editing rates at endogenous genomic loci, cells were cultured 3 days following transfection, after which the media was removed, the cells were harvested, and genomic DNA was isolated using QIAamp DNA mini kit (QIAGEN) according to the manufacturer’s instructions. The editing rates were then determined by targeted amplicon deep sequencing.

### Electroporation of iMyoblasts and Fibroblast cells

PEmax mRNA mixed with synthetic pegRNA/sgRNA mixtures or RNPs programmed with synthetic pegRNA/sgRNA were delivered by electroporation using the NEON Nucleofection System 10 μL kit (Thermo Fisher Scientific). For PEmax mRNA-based editing experiments in fibroblasts, 100k cells were pelleted at 300 g for 5 min and resuspended in 10 μL NEON Buffer R containing 1 μg PEmax mRNA, 100 pmol synthetic pegRNA (IDT), 15 pmol synthetic sgRNA (IDT, for PE3 approaches; Supplementary Table) and 500 ng of VPX/VPX^Q76A^ mRNA (for VPX mRNA co-delivery). The NEON Nucleofection System (Invitrogen) was used for electroporation with 10 μL tips (HEK293T: 1150v, 20ms, 2 pulses; U2OS: 1200v, 20ms, 2 pulses). For editing experiments in iMyoblasts, Limb girdle muscular dystrophy 2G (LGMD2G) iMyoblasts were differentiated from LGMD2G iPSCs and maintained as previously described^26,49,50^. For mRNA-based editing experiments, 200,000 iMyoblasts cells were resuspended into 10 µl of NEON Buffer R, mixed with 1μg PEmax variant mRNA, 100 pmol synthetic pegRNA (IDT) and 50pmol sgRNA (IDT), and then electroporated using NEON Nucleofection System 10 μL kit (Thermo Fisher Scientific) as follows: pulse voltage 1,500 V, pulse width 20 ms, pulse number 1. After electroporation, the cells were plated into a 6-well plate with HMP growth medium. For RNP-based editing experiments, 50 pmol of PEmax protein was incubated with 200 pmol of synthetic pegRNA (IDT) and 15 pmol of sgRNA (IDT) in R buffer to a total volume of 10 μL for 15 min at room temperature for complex formation. 200k cells were electroporated with 10 μL of PEmax RNP complex using the same electroporation and culture conditions described above for mRNA delivery. gDNA was isolated 3 days after electroporation from each group and stored at −80°C for Illumina library preparation.

### Prime editing experiments in human primary CD4^+^ T cells

Leukopaks were obtained from anonymous, healthy blood donors (New York Biologics, Southhampton, New York); these experiments were reviewed by the University of Massachusetts Medical School Institutional Review Board and declared to be non-human subjects research, according to National Institutes of Health (NIH) guidelines (http://grants.nih.gov/grants/policy/hs/faqs_aps_definitions.htm). To generate primary CD4+ T cells, peripheral blood mononuclear cells (PBMCs) were isolated from human donor leukopaks by gradient centrifugation on lymphoprep (cat#07861, Stemcell Technologies). Thereafter, PBMCs were depleted of CD14 mononuclear cells using anti-CD14 microbead antibodies (cat#130-050-201, Miltenyi Biotec) and the flowthrough was enriched for CD4+ T cells by positive selection using anti-CD4 microbead antibodies (cat#130-045-101, Miltenyi Biotec). CD4+ T cell enrichment was confirmed by determining the percentage of CD3+/CD4+ cells via flow cytometry. Isolated CD4+ T cells were cultured in complete RPMI-IL2 media (RPMI-1640 media (cat# 11875093, Thermofisher Scientific) supplemented with 10% heat-inactivated Cosmic Calf Serum (cat#SH30087.03, GE Life Sciences), 25 mM HEPES pH 7.2 (cat#25-060-CI, Corning), 20 mM GlutaMAX (cat#3505-061, Gibco), 1 mM Sodium pyruvate (cat#25-000-CI, Corning), 1X MEM non-essential amino acids (cat#25-025-CI, Corning), 1% penicillin-streptomycin (cat#15140-122, Gibco), and 1:2000 human interleukin-2 (made in-house from IL-2 expressing cell line). 3 days prior to electroporation, primary CD4+ T cells were activated with anti-CD3/CD28 antibodies (cat#10971, Stemcell Technologies). 1e6 activated primary CD4+ T cells were electroporated with 1μg PEmax variant mRNA, 100pmol synthetic pegRNA (IDT) and 50pmol sgRNA (IDT) using the P3 primary cell nucleofector kit (cat#V4XP-3032, Lonza Biosciences) and program EH-115 on an Amaxa 4D-Nucleofector. For the resting CD4+ T cell experiments, 1e6 cells were electroporated using the P3 primary cell nucleofector kit and program FI-115 on the Amaxa 4D-Nucleofector immediately after isolation. CD4+ T cells were allowed to recover in RPMI-IL2 media for 72 hours post nucleofection before genomic extraction using the Qiagen QiAamp DNA Blood Mini Kit (cat#51104, Qiagen). For VPX co-delivery experiments, 0.5 - 4 μg of VPX protein (ab267924, Abcam) or 500 ng of VPX/VPX^Q76A^ mRNA was added to the electroporation mixture described above.

### Zebrafish prime editing experiments

Zebrafish were maintained and bred according to standard protocols set by University of Massachusetts Chan Medical School Institutional Animal Care and Use Committee (IACUC). Zebrafish embryos obtained from EK (WT) wild-type in-crosses were used for one cell-stage microinjections of PE RNPs. Prior to injections the *tek* target sequence was verified by Sanger sequencing. For 1xRNP, 12 mM pegRNA (synthesized by IDT) and 6 mM PE protein were combined in nuclease-free water. For 2x and 4X RNP, the amount of pegRNA and protein used were scaled up 2-fold and 4-fold respectively. Complexes were incubated at room temperature for 5 minutes and then 2 nl was injected into single-cell embryos. Injected embryos were incubated at 28.5 °C overnight. Twenty-four hours post injection embryos were assessed for toxicity and genomic DNA was extracted from 20 normally developing embryos using the Qiagen DNeasy Blood and Tissue kit (Qiagen). Injections were performed in three independent replicates.

### *In vivo* delivery of PEmax mRNA/pegRNA/sgRNA delivery for *Pcsk9* editing in mice

PEmax and PEmax** mRNA were *in vitro* transcribed and purified as described above. For *in vivo* mouse experiments, an additional round of purification using cellulose was performed to remove dsRNA contaminants^51^. The purified PEmax/PEmax** mRNA (40mg) was mixed with Pcsk9 synthetic pegRNA (15mg, IDT) and nicking sgRNA (5mg, IDT) in 10mM citrate buffer. This RNA mix constituted the aqueous component for forming RNA-loaded LNPs by microfluidic mixing using the NanoAssemblr Ignite (Precision Nanosystems) with a 3:1 aqueous:ethanol flow rate ratio and a total flow rate of 12mL/min. The ethanol component consisted of a lipid mix of D-Lin-MC3-DMA (MedChem Express: HY-112251), DSPC (Sigma-Aldrich: P1138), cholesterol (Sigma-Aldrich: C8667), and DMG-PEG at a 50:10:38.5:1.5 molar ratio^52^. A lipid to nucleic acid weight ratio of 20 was used to determine mixing volume. After formulation, LNPs were dialyzed (Sigma-Aldrich: PURX35050) overnight at 4°C in 1xDPBS (Thermo: 14190250), sterile filtered (Thermo: 723-2520), and concentrated (Millipore: UFC8100) to 150ml per dose. Quality control was performed by dynamic light scattering (Malvern ZetaSizer) to assess particle diameter & RiboGreen (Invitrogen: R11490) quantification of RNA encapsulation efficiency.

Mouse protocols were approved by the UMass Chan Medical School IACUC. LNPs containing mRNA were delivered by retro-orbital injection into B6 wild type female mice (three doses spaced one week apart). Fresh LNPs were formulated for each dose. Mice were sacrificed one week after the final dose and the liver was harvested. Three liver punches from each mouse liver (one from each lobe) were taken for gDNA extraction. The editing rates were then determined by targeted amplicon deep sequencing.

### Targeted amplicon deep sequencing to assess editing rates

Genomic DNA was isolated for prime editing analysis from treated cells or zebrafish embryos. Genomic loci spanning each target site were PCR amplified with locus-specific primers carrying tails complementary to the Truseq adapters. 200 ng of genomic DNA was used for the 1^st^ PCR using Phusion master mix (Thermo) with locus specific primers that contain tails. PCR products from the 1^st^ PCR were used for the 2^nd^ PCR with i5 primers and i7 primers to complete the adaptors and include the i5 and i7 indices. All primers used for the amplicon sequencing are listed in the supplementary table. PCR products were purified with Ampure beads (0.9X reaction volume), eluted with 25ul of TE buffer, and quantified by Qubit. Equal molar ratios of each amplicon were pooled and sequenced using Illumina Miniseq. Amplicon sequencing data was analyzed with CRISPResso (https://crispresso.pinellolab.partners.org/)^53^. Briefly, demultiplexing and base calling were both performed using bcl2fastq Conversion Software v2.18 (Illumina, Inc.), allowing 0 barcode mismatches with a minimum trimmed read length of 75. Alignment of sequencing reads to each amplicon sequence was performed using CRISPResso2. For single base substitution, samples were analyzed by CRISPResso2 in regular batch mode using the following parameters: ‘‘-q 30’’, “--discard_indel_reads TRUE’’, and “-qwc (or -- quantification_window_coordinates)”. The qwc value specifies the quantification window for indel analysis, including the entire sequence between pegRNA and nicking sgRNA-directed Cas9 nicking sites (when TWIN-PE or PE3 is employed), as well as an additional 10 bp beyond both cut sites. Intended PE editing efficiency was calculated as the percentage of reads with the desired edit without indels (‘‘-discard_indel_reads TRUE.’’ mode) out of the total number of reads ((number of desired edit-containing reads) / (number of reference-aligned reads)). Unintended editing frequency was calculated as the number of discarded reads divided by the total number of reads ((number of indel-containing reads) / (number of reference-aligned reads)). When the desired prime editing outcome comprises multiple base changes or an insertion or deletion, samples were analyzed by CRISPResso2 in regular batch mode using the following parameters: ‘‘-q 30’’, “--discard_indel_reads TRUE’’, “-qwc” and ‘‘--expected_hdr_amplicon_seq’’ providing the desired edited amplicon sequence for each target site. For these types of edits, the intend edit was calculated by (the number of HDR-aligned reads) / (number of reference-aligned reads). Unintended edits were calculated as the number of discarded reads divided by the total number of reads ((number of reads ‘‘Discarded’’ reads from the reference-aligned sequences + number of the ‘‘Discarded’’ reads from the HDR-aligned sequences)/(number of reference-aligned reads)). All these values can be found in the CRISPResso2 output file ‘‘CRISPResso_quantification_of_editing_frequency.txt’’. The analysis of scaffold sequences incorporated within the unintended edited sequence population was determined using the scaffold sequence analysis pipeline previously described^12^, where incorporation of at least 2 bases of the scaffold sequence were required to assign the read as containing a scaffold insertion.

### Data availability

A Reporting Summary for this article is available as a supplementary information file. Plasmids for mRNA *in vitro* transcription and protein expression of the new prime editor variants have been deposited to Addgene for distribution. Illumina Sequencing data have been submitted to the Sequence Read Archive, and datasets are available under BioProject accession number PRJNA1024467. Source data are provided with this paper.

---

**Extended Data Fig. 1:**
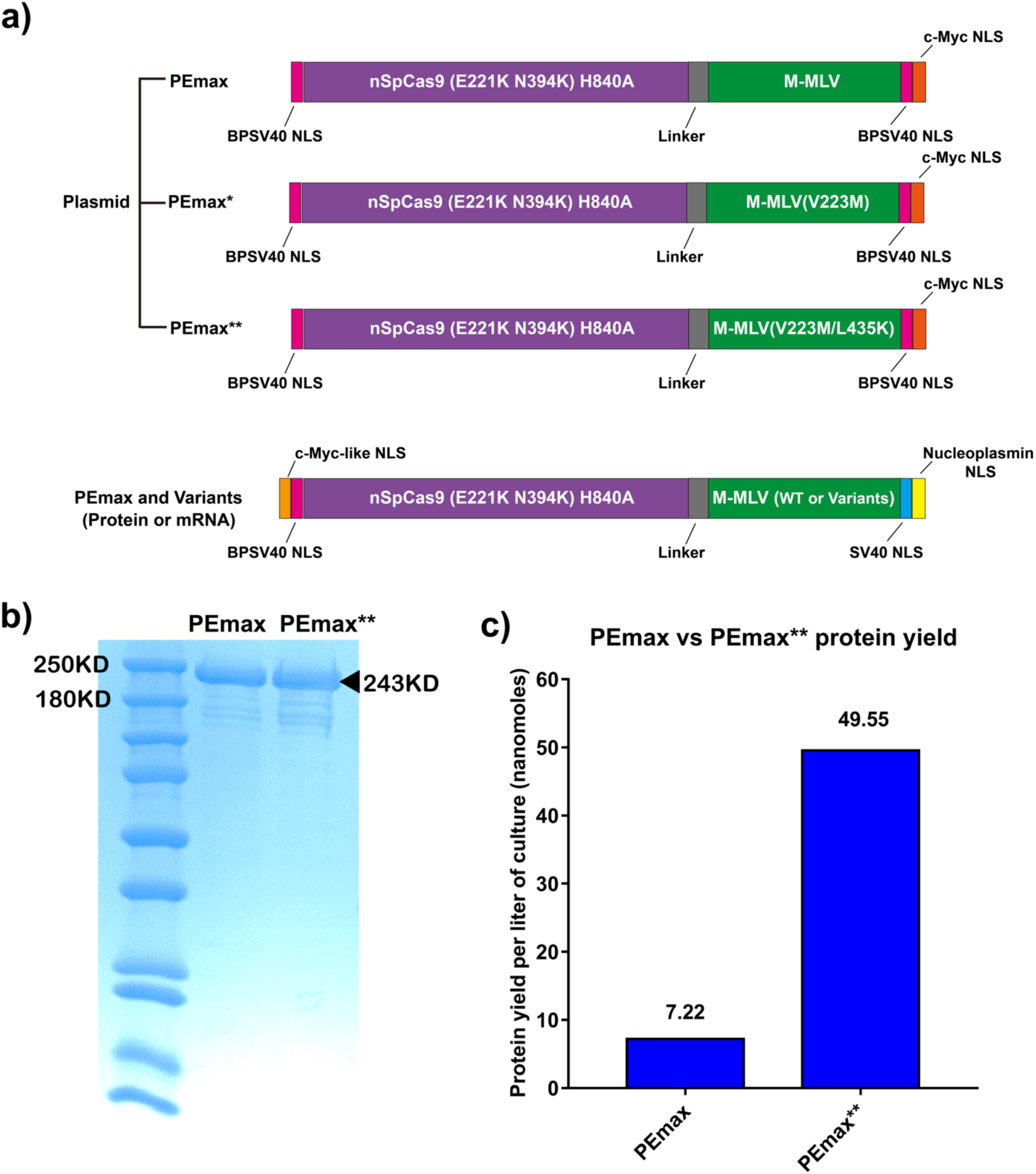
PEmax protein expression constructs and their purification. **(a)** Architecture of the PEmax, PEmax* and PEmax** mammalian expression plasmid, mRNA and protein constructs. Note that PEmax variants encoded by mRNA and protein constructs contain four nuclear localization signal (NLS) sequences^6^ whereas the mammalian expression plasmid encoded PEmax variants contain three NLSs.^2^ **(b)** SDS-PAGE analysis of purified C-terminally His-tagged PEmax and PEmax** protein. Black arrow indicates the desired protein product. **(c)** Protein yield per liter of bacterial culture for PEmax and PEmax**.

**Extended Data Fig. 2:**
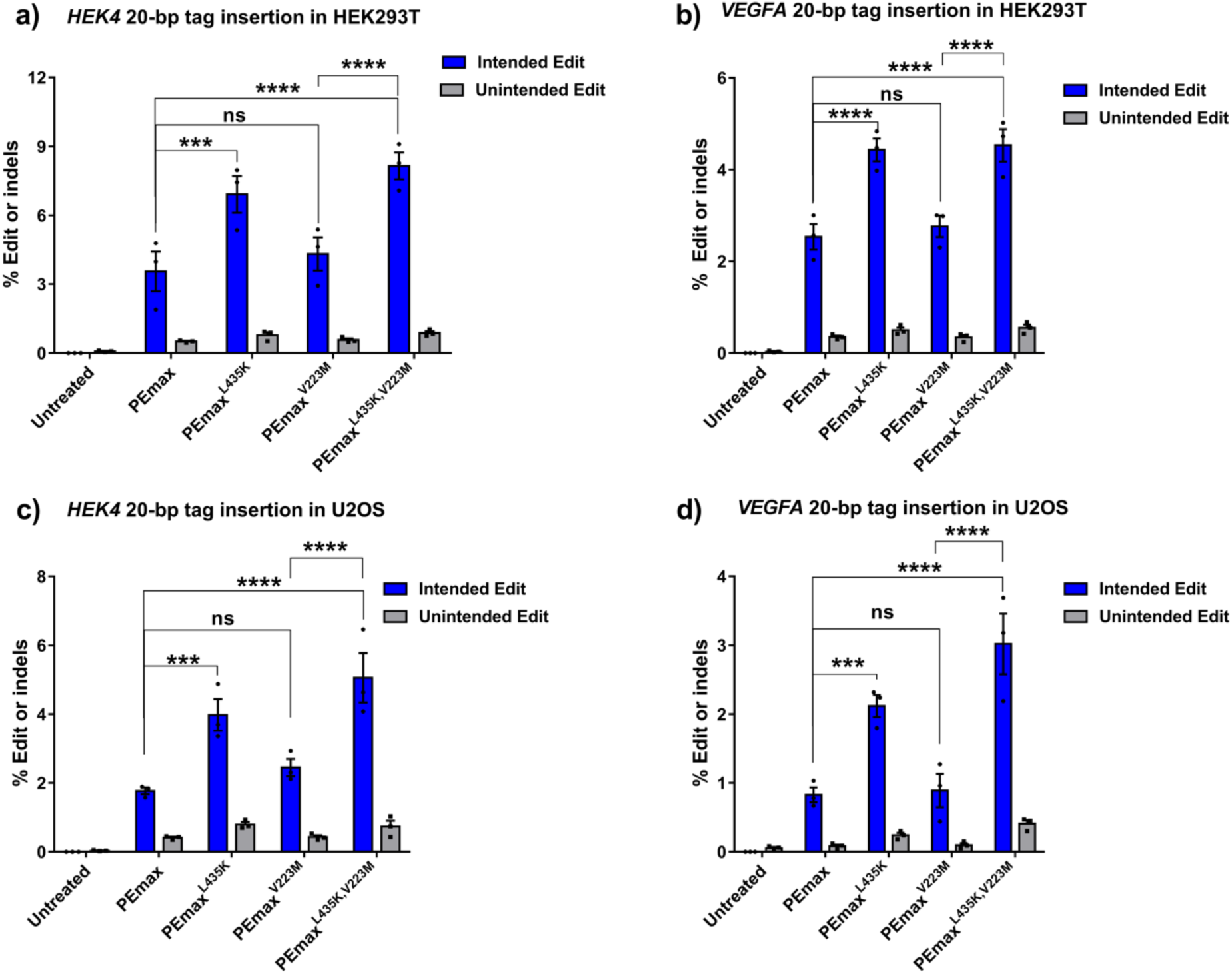
MMLV-RT^L435K^ improves prime editing rates in transformed cell lines. PEmax, PEmax (L435K), PEmax* (V223M) and PEmax** (V223M & L435K) mediated 20 bp tag insertions at **(a)** *HEK4* and **(b)** *VEGFA* in HEK293T cells (n=3 independent replicates) and **(c)** *HEK4* and **(d)** *VEGFA* in U2OS cells (n=3 independent replicates). Expression plasmids were delivered by transient transfection. Two-way ANOVA was used to compare the intended edit and unintended edits across all the groups for each graph, and PEmax was used as a control column for multiple comparisons; ns indicates P > 0.05, ** indicates P ≤0.01, *** indicates P ≤0.001 and **** indicates P ≤0.0001 (also see Supplementary table).

**Extended Data Fig. 3:**
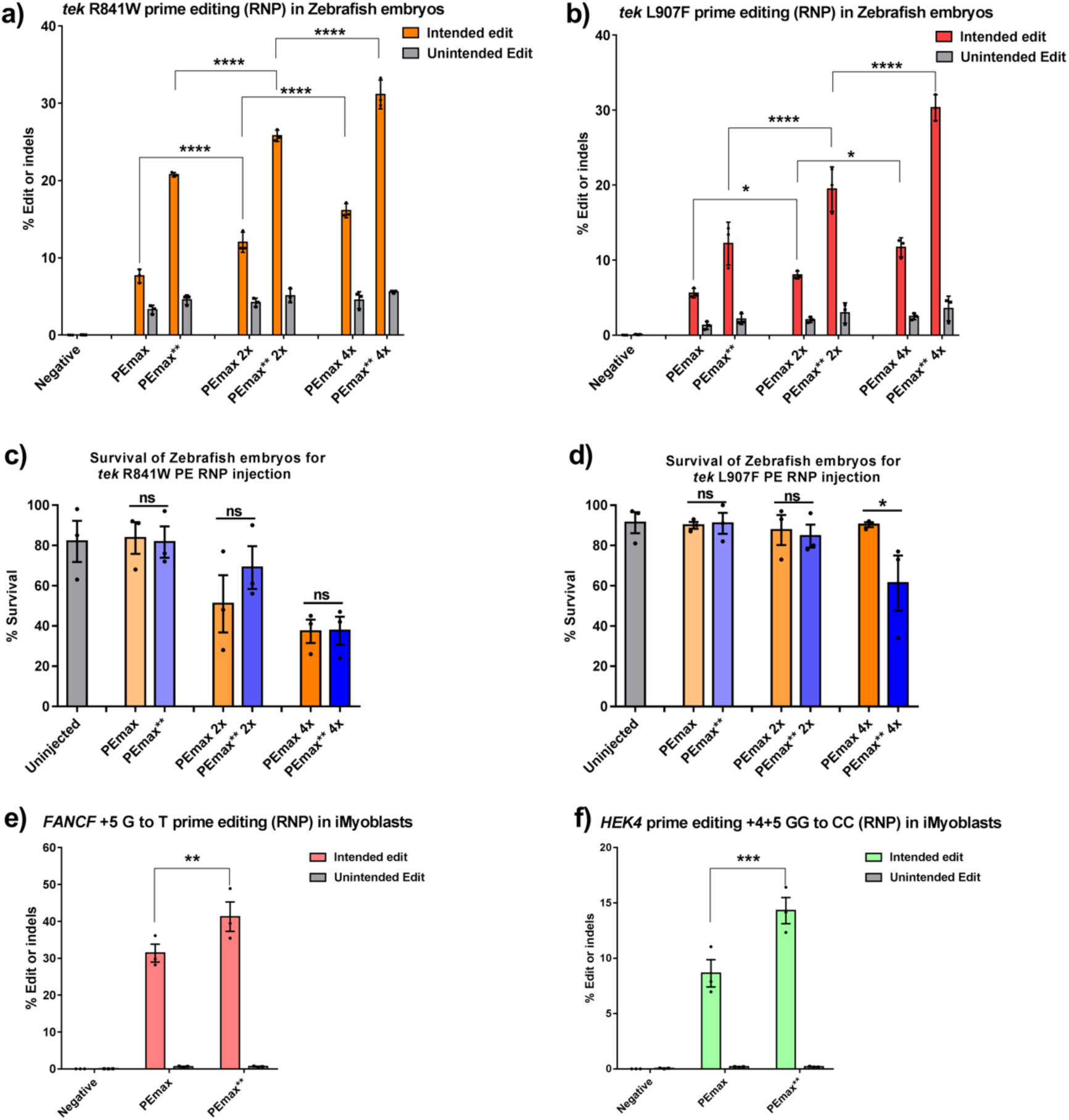
PEmax** protein improves editing rates in zebrafish and induced myoblasts. Comparison of PEmax and PEmax** RNP editing efficiency at various concentrations for introducing **(a)** R841W and **(b)** L907F at the *tek* locus in zebrafish embryos (n=3 independent replicates). One-way ANOVA was used to compare the intended edit across all the groups for each graph; * indicates P < 0.05, **** indicates P ≤0.0001 (also see Supplementary table). Comparison of zebrafish embryo survival rate 24 hours post RNP injection for different treatment groups for **(c)** R841W and **(d)** L907F at the *tek* lokus. One-way ANOVA was used to compare the intended edit across all the groups for each graph; ns indicates P > 0.05, * indicates P < 0.05 (also see Supplementary table). Comparison of PEmax and PEmax** RNP editing efficiency in mediating PE3 editing at **(e)** *FANCF* (+5 G to T) and (**f**) HEK4 (+4+5 GG to CC) in induced myoblasts (n=3 independent replicates). Two-way ANOVA was used to compare the intended edit and unintended edits across all the groups for each graph, and PEmax was used as a control column for comparisons; ** indicates P ≤0.01, *** indicates P ≤0.001 (also see Supplementary table).

**Extended Data Fig. 4:**
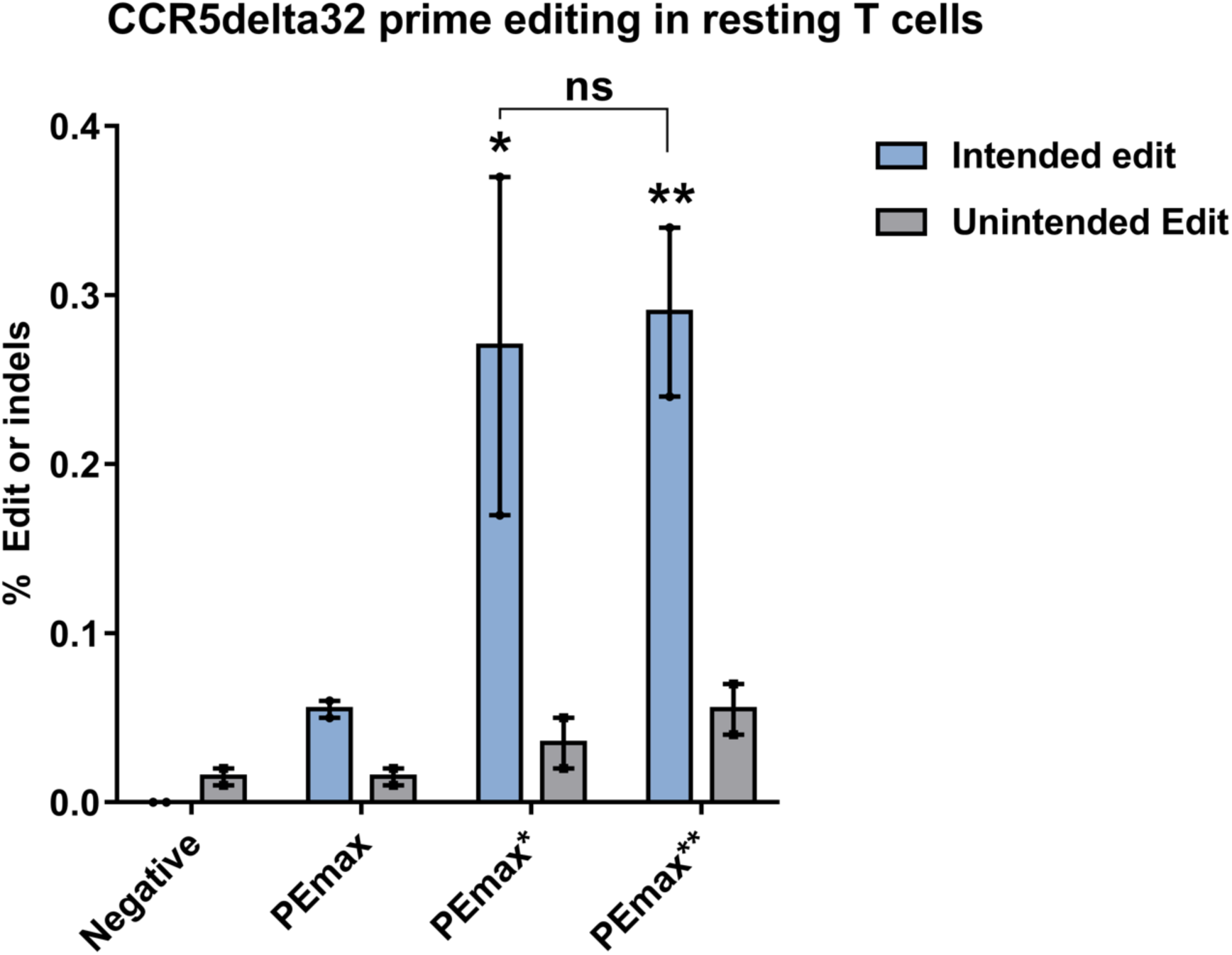
Comparison of PEmax variants in resting T cells. PEmax, PEmax* and PEmax** mRNA mediated PE3 editing to install *CCR5*^delta32^ HIV-1 resistance mutation in resting human primary T cells (n=2 different donors). Two-way ANOVA was used to compare the intended edit and unintended edits across all the groups for each graph, and PEmax was used as a control column for multiple comparisons; ns indicates P > 0.05, * indicates P < 0.05, ** indicates P ≤0.01 (also see Supplementary table).

**Extended Data Fig. 5:**
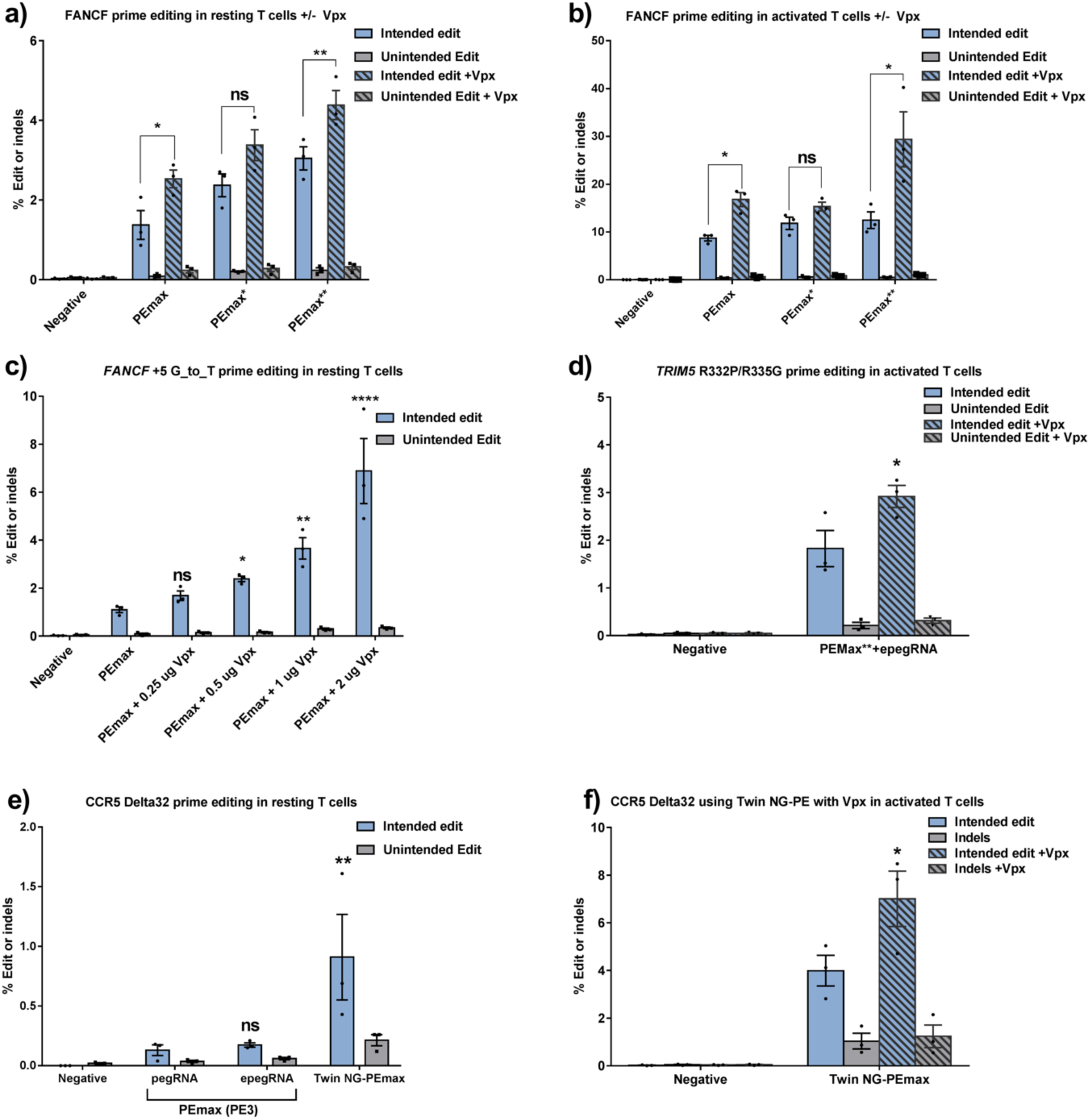
Co-delivery of VPX protein improves prime editing rates. PEmax, PEmax*, and PEmax** mRNA mediated PE3 editing at *FANCF* (+5 G to T) in **(a)** resting and **(b)** activated human primary T cells with co-delivery of synthetic pegRNA/sgRNA and 0.5 μg VPX protein (n=3 different donors). Two-way ANOVA was used to compare the intended edit and unintended edits across all the groups for each graph, and the no VPX group was used as a control column for multiple comparisons; ns indicates P > 0.05, * indicates P ≤0.05, ** indicates P ≤0.01 (also see Supplementary table). **(c)** PEmax-mRNA mediated PE3 editing at *FANCF* (+5 G to T) in resting human primary T cells with co-delivery of synthetic pegRNA/sgRNA and increasing amounts of VPX protein (n=3 different donors). Two-way ANOVA was used to compare the intended edit and unintended edits across all the groups for each graph, and the no VPX group was used as a control column for multiple comparisons; ns indicates P > 0.05, * indicates P ≤0.05, ** indicates P ≤0.01 and **** indicates P ≤0.0001 (also see Supplementary table). (**d**) PE3 editing to install *TRIM5ɑ*^R332P,R335G^ HIV-1 resistance-associated mutation in primary human T cells with co-delivery of 2 μg VPX protein with PEmax** and a synthetic epegRNA (n=3 different donors). Two-way ANOVA was used to compare the intended edit and Indels between two groups, and the PEmax**+epegRNA without VPX group was used as a control column for multiple comparisons; * indicates P ≤0.05 (also see Supplementary table). **(e)** Comparison of PEmax-mediated PE3 editing (pegRNA/epegRNA) and a Cas9-NG PEmax based TWIN-PE approach for installing *CCR5^delta^*^32^ HIV-1 resistance mutation in resting human primary T cells. Two-way ANOVA was used to compare the intended edit and Indels across all the groups, and the PEmax**+pegRNA group was used as a control column for multiple comparisons; ns indicates P > 0.05, ** indicates P < 0.01 (also see Supplementary table). (**f**) Twin-PE editing to install *CCR5*^delta32^ HIV-1 resistance mutation in human primary T cells with co-delivery of 2 μg VPX protein and Cas9-NG PEmax** (n=3 different donors). Two-way ANOVA was used to compare the intended edit and Indels across all the groups, and the no VPX group was used as a control column for multiple comparisons; * indicates P < 0.05 (also see Supplementary table).

**Extended Data Fig. 6:**
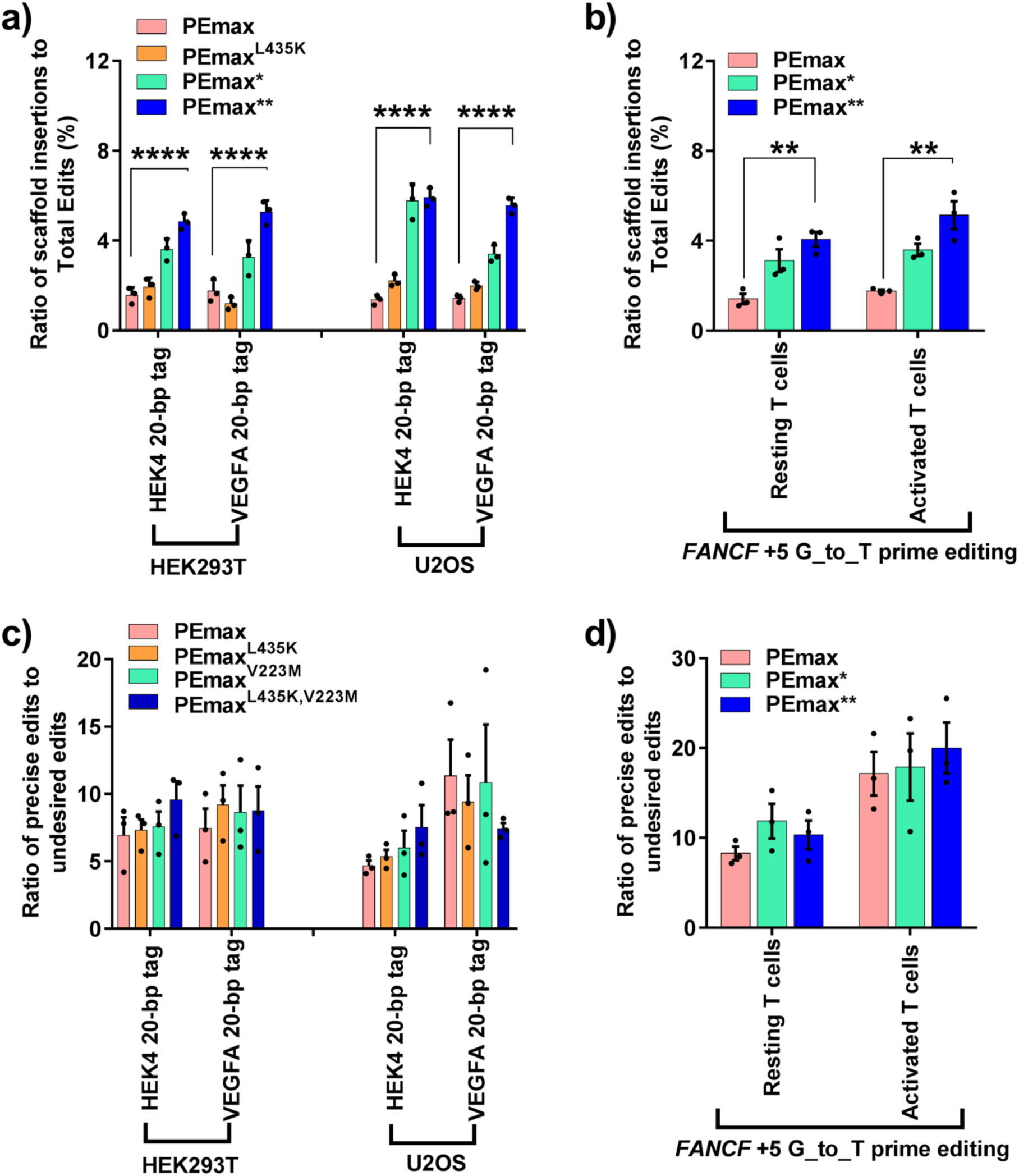
Analysis of undesired editing outcomes after prime editing with PEmax or PEmax variants. **(a, b)** Percentage of sequencing reads containing a pegRNA scaffold sequence insertion within the population of total edited sequences (both precise and undesired edits) after prime editing using PEmax and PEmax variants in different cell types. **(a)** Scaffold sequence insertions for prime editing by PEmax, PEmax^L435K^, PEmax* and PEmax** at HEK4 and VEGFA target sites in HEK293T cells and U2OS cells from Extended Data Fig. 2. (b) Scaffold sequence insertions for prime editing by PEmax, PEmax* and PEmax** without VPX at the FANCF target site in resting and activated T cells from Extended Data Fig. 5a,b. One-way ANOVA was used to compare the ratio of scaffold insertion across each graph, ** indicates P ≤0.01, **** indicates P ≤0.0001 (also see Supplementary table). **(c, d)** Ratio of precise edits to undesired edits in **(a, b),** no significant difference was detected across these groups (also see Supplementary table). Bars reflect the mean of n = 3 independent replicates. Dots show individual replicate values.

**Extended Data Fig. 7:**
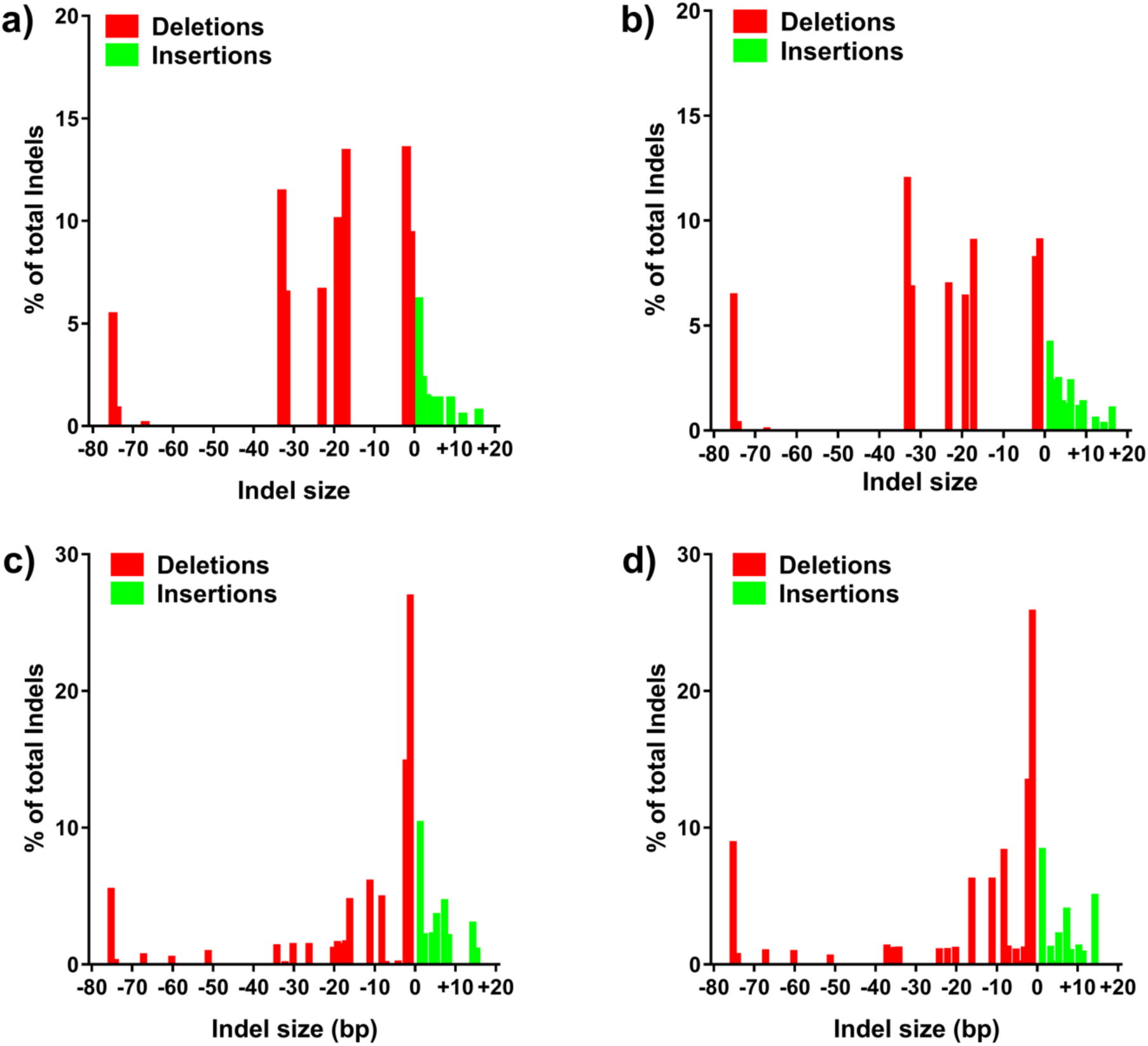
Analysis of Indel distribution after prime editing with or without VPX. **(a-d)** Histograms of indel size distribution for PEmax** mRNA-mediated PE3 editing with co-delivery of synthetic pegRNA/sgRNA at *FANCF* (+5 G to T). Prime editing in resting human primary T cells without **(a)** or with **(b)** 0.5 μg VPX protein, and prime editing in activated human primary T cells without **(c)** and with **(d)** 0.5 μg VPX protein (see Extended Data Figure 5a,b for primary editing data, n=3 different donors).

