## Supplementary Data and Figures for "Addressing the dNTP bottleneck restricting prime editing activity"

### Supplementary Note

#### Rationale behind choosing particular MMLV-RT mutants for generating prime editor variants

In our previous study, we observed the propensity for protein aggregation at various steps during purification of the prime editor protein[1]. The poor solubility of the prime editor protein is likely a function of the MMLV-RT domain, as purified Cas9 protein does not display these adverse solubility characteristics as an independent protein[1,2]. A 24 amino acid deletion at the N terminal region of MMLV-RT or L435K substitution has been shown to increase solubility of the protein[3]. However, the 24 amino acid truncation has been shown to decrease prime editing rates[4]. Therefore, we decided to evaluate the L435K mutation, which is a solvent exposed residue among a hydrophobic stretch of amino acids in the connector domain of MMLV-RT, in the context of prime editing. Additionally, since dNTP levels are typically much lower in non-dividing cells than dividing cells[5] and MMLV-RT has reduced RNA dependent DNA polymerization activity under low dNTP concentrations[6], we hypothesized that cellular dNTP levels could be a factor limiting prime editing efficiency. Q221R and V223A/M substitutions have been shown to increase the dNTP affinity[7]. However, Q221R demonstrates a reduced fidelity[7] and therefore we avoided testing this mutation. We chose the V223M substitution since it alters the conserved Y**V**DD motif in MMLV-RT to Y**M**DD, which changes the active site from an onco-retroviral RT to a more efficient lentiviral-like RT, reducing its $K_m$ for dNTPs by 2 to 4-fold[7–9].


1. Ponnienselvan, K. *et al.* Reducing the inherent auto-inhibitory interaction within the pegRNA enhances prime editing efficiency. *Nucleic Acids Res.* **51**, 6966–6980 (2023).
2. Wu, Y. *et al.* Highly efficient therapeutic gene editing of human hematopoietic stem cells. *Nat. Med.* **25**, 776–783 (2019).
3. Das, D. & Georgiadis, M. M. A directed approach to improving the solubility of Moloney murine leukemia virus reverse transcriptase. *Protein Sci.* **10**, 1936–1941 (2001).
4. Grünewald, J. *et al.* Engineered CRISPR prime editors with compact, untethered reverse

transcriptases. *Nat. Biotechnol.* **41**, 337–343 (2022).

**Supplementary figure 1**

PEmax** sequence –mRNA expression vector

c-Myc_like_NLS-BPSV40_NLS-nSpCas9 (E221K N394K) H840A-linker-MMLV-RT (V223M,L435K)-SV40_NLS-Nucleoplasmin_NLS-6HIS-tag

```
MPAAKKKKLDGSVDKRTADGSEFESPKKKRKVDKKYSIGLDIGTNSVGWAVITDEYKVPSKKFKVLGNTD
RHSIKKNLIGALLFDSGETAEATRLKRTARRRYTRRKNRICYLQEIFSNEMAKVDDSFFHRLEESFLVEE
DKKHERHPIFGNIVDEVAYHEKYPTIYHLRKKLVDSTDKADLRLIYLALAHMIKFRGHFLIEGDLNPDNS
DVDKLFIQLVQTYNQLFEENPINASGVDAKAILSARLSKSRKLENLIAQLPGEKKNGLFGNLIALSLGLT
PNFKSNFDLAEDAKLQLSKDTYDDDLDNLLAQIGDQYADLFLAAKNLSDAILLSDILRVNTEITKAPLSA
SMIKRYDEHHQDLTLLKALVRQQLPEKYKEIFFDQSKNGYAGYIDGGASQEEFYKFIKPILEKMDGTEEL
LVKLKREDLLRKQRTFDNGSIPHQIHLGELHAILRRQEDFYPFLKDNREKIEKILTFRIPYYVGPLARGN
SRFAWMTRKSEETITPWNFEEVVDKGASAQSFIERMTNFDKNLPNEKVLPKHSLLYEYFTVYNELTKVKY
VTEGMRKPAFLSGEQKKAIVDLLFKTNRKVTVKQLKEDYFKKIECFDSVEISGVEDRFNASLGTYHDLLK
IIKDKDFLDNEENEDILEDIVLTLTLFEDREMIEERLKTYAHLFDDKVMKQLKRRRYTGWGRLSRKLING
IRDKQSGKTILDFLKSDGFANRNFMQLIHDDSLTFKEDIQKAQVSGQGDSLHEHIANLAGSPAIKKGILQ
TVKVVDELVKVMGRHKPENIVIEMARENQTTQKGQKNSRERMKRIEEGIKELGSQILKEHPVENTQLQNE
KLYLYYLQNGRDMYVDQELDINRLSDYDVDAIVPQSFLKDDSIDNKVLTRSDKNRGKSDNVPSEEVVKKM
```

KNYWRQLLNAKLITQRKFDNLTKAERGGLSELDKAGFIKRQLVETRQITKHVAQILDSRMNTKYDENDKL
IREVKVITLKSKLVSDFRKDFQFYKVREINNYHHAHDAYLNAVVGTALIKKYPKLESEFVYGDYKVYDVR
KMIAKSEQEIGKATAKYFFYSNIMNFFKTEITLANGEIRKRPLIETNGETGEIVWDKGRDFATVRKVLSM
PQVNIVKKTEVQTGGFSKESILPKRNSDKLIARKKDWDPKKYGGFDSPTVAYSVLVVAKVEKGKSKKLKS
VKELLGITIMERSSFEKNPIDFLEAKGYKEVKKDLIIKLPKYSLFELENGRKRMLASAGELQKGNELALP
SKYVNFLYLASHYEKLKGSPEDNEQKQLFVEQHKHYLDEIIEQISEFSKRVILADANLDKVLSAYNKHRD
KPIREQAENIIHLFTLTNLGAPAAFKYFDTTIDRKRYTSTKEVLDATLIHQSITGLYETRIDLSQLGGDS
GGSSGGSKRTAGSYPYDVPDYADGSEFESPKKKRKVSGGSSGGSTLNIEDEYRLHETSKEPDVSLGSTWL
SDFPQAWAETGGMGLAVRQAPLIIPLKATSTPVSIKQYPMSQEARLGIKPHIQRLLDQGILVPCQSPWNT
PLLPVKKPGTNDYRPVQDLREVNKRVEDIHPTVPNPYNLLSGLPPSHQWYTVLDLKDAFFCLRLHPTSQP
LFAFEWRDPEMGISGQLTWTRLPQGFKNSPTLFNEALHRDLADFRIQHPDLILLQY**M**DDLLLAATSELDC
QQGTRALLQTLGNLGYRASAKKAQICQKQVKYLGYLLKEGQRWLTEARKETVMGQPTPKTPRQLREFLGK
AGFCRLFIPGFAEMAAPLYPLTKPGTLFNWGPDQQKAYQEIKQALLTAPALGLPDLTKPFELFVDEKQGY
AKGVLTQKLGPWRRPVAYLSKKLDPVAAGWPPCLRMVAAIAVLTKDAGKLTMGQPLVI**K**APHAVEALVKQ
PPDRWLSNARMTHYQALLLDTDRVQFGPVVALNPATLLPLPEEGLQHNCLDILAEAHGTRPDLTDQPLPD
ADHTWYTDGSSLLQEGQRKAGAAVTTETEVIWAKALPAGTSAQRAELIALTQALKMAEGKKLNVYTDSRY
AFATAHIHGEIYRRRGWLTSEGKEIKNKDEILALLKALFLPKRLSIIHCPGHQKGHSAEARGNRMADQAA
RKAAITETPDTSTLLIENSSPSGGSTGGGPGGGAAAGSGSPKKKRKVGSGSKRPAATKKAGQAKKKKLEH
HHHHH
